# RADpainter and fineRADstructure: population inference from RADseq data

**DOI:** 10.1101/057711

**Authors:** Milan Malinsky, Emiliano Trucchi, Daniel John Lawson, Daniel Falush

## Abstract

Powerful approaches to inferring recent or current population structure based on nearest neighbour haplotype ‘coancestry’ have so far been inaccessible to users without high quality genome-wide haplotype data. With a boom in non-model organism genomics, there is a pressing need to bring these methods to communities without access to such data. Here we present `RADpainter`, a new program designed to infer the coancestry matrix from restriction-site-associated DNA sequencing (RADseq) data. We combine this program together with a previously published MCMC clustering algorithm into `fineRADstructure` - a complete, easy to use, and fast population inference package for RADseq data (https://github.com/millanek/fineRADstructure). Finally, with two example datasets, we illustrate its use, benefits, and robustness to missing RAD alleles in double digest RAD sequencing.

## Introduction

Understanding of shared ancestry in genetic datasets is often key to their interpretation. The `chromoPainter/fineSTRUCTURE` package (Lawson et al. 2012) represents a powerful model-based approach to investigating population structure using genetic data. It offers especially high resolution in inference of recent shared ancestry, as shown for example in the investigations of worldwide human population history (Hellenthal et al. 2014) and of genetic structure of the British population (Leslie et al. 2015). Further advantages, when compared with other model-based methods, such as STRUCTURE (Pritchard et al. 2000) and ADMIXTURE (Alexander et al. 2009), include the ability to deal with a very large number of populations, explore relationships between them, and to quantify ancestry sources in each population.

The high resolution of `chromoPainter/fineSTRUCTURE` and related methods derives from utilizing haplotype linkage information and from focusing on the most recent coalescence (common ancestry) among the sampled individuals. This approach derives a ‘coancestry matrix’, a summary of nearest neighbour haplotype relationships in the dataset; i.e. of the cases where pairs of individuals had the most similar haplotypes one to another. However, the existing pipeline for coancestry matrix inference was designed for large scale human genetic SNP datasets, where chromosomal location of the markers are known, haplotypes are typically assumed to be correctly phased (although it is possible to perform the analysis without this assumption), and missing data also needs to have been imputed. Therefore, these methods have so far been generally inaccessible for investigations beyond model organisms.

With no requirements for prior genomic information (e.g. no need for a reference genome) and relatively low cost, RADseq data are fuelling a boom in ecological and evolutionary genomics, especially for non-model organisms (Andrews et al. 2016), where questions on population structure and relative ancestry are among the most commonly asked. Therefore, we have developed `RADpainter`, a simple method designed to infer the coancestry matrix from RADseq data. `RADpainter` is designed to take full advantage of the sequence of all the SNPs from each RAD locus, finding (one or more) closest relatives for each allele. The information about the nearest neighbours of each individual is then summed up into the coancestry similarity matrix. We package `RADpainter` together with the `fineSTRUCTURE` Markov chain Monte Carlo (MCMC) clustering algorithm into an easy to use population inference package for RADseq data called `fineRADstructure`.

## New Approaches

Briefly, the coancestry matrix is calculated as follows: for each RAD locus and each individual (a recipient), we calculate the number of sequence differences (i.e. SNPs) between that individual’s allele(s) and the alleles in all other individuals (potential donors). The closest relative (donor) for each allele is its nearest neighbour allele, i.e. the allele with the least number of differences. In the case of multiple equally distant nearest neighbours, an equal proportion of coancestry is assigned to each ‘donor’. Finally, we sum these local coancestry values across all loci to obtain the coancestry matrix for the full dataset. A basic outline of the coancestry estimation procedure for haploid individuals is shown in Algorithm 1.

**Algorithm 1.**
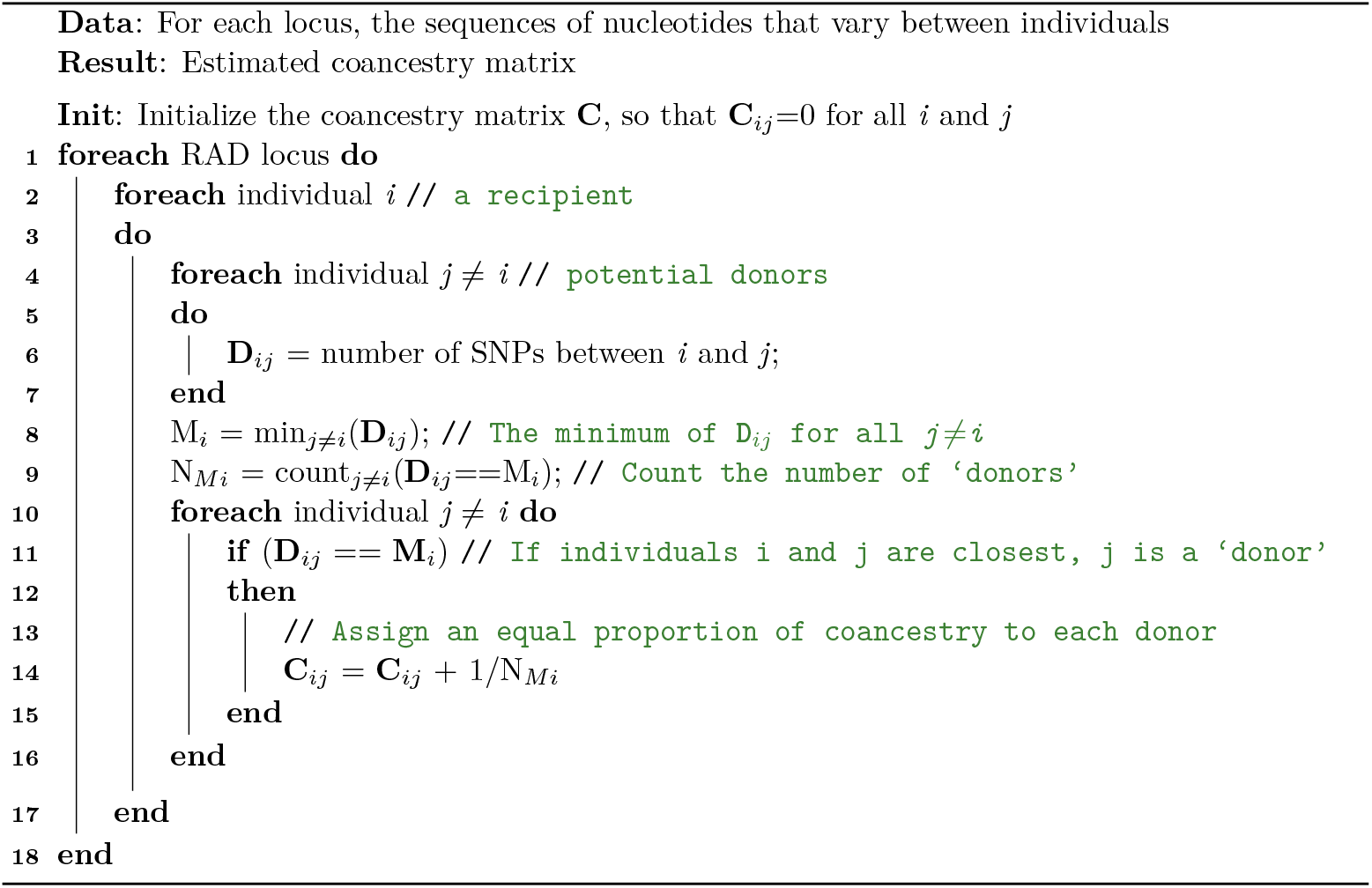
Coancestry estimation: a simple case with haploid individuals and without missing data.

Differences in ploidy are handled by averaging coancestry across alleles in the same individual. Figure 1 provides a further illustration of the method, showing co-ancestry matrix calculation at a single locus for four diploid individuals. However, we have implemented `RADpainter` in a way to handle arbitrary number of alleles per individual (i.e. any ploidy levels), depending on the input; ploidy can even vary across individuals in a single analysis. We expect these features to be of particular use to the plant research community.

**Figure 1.**
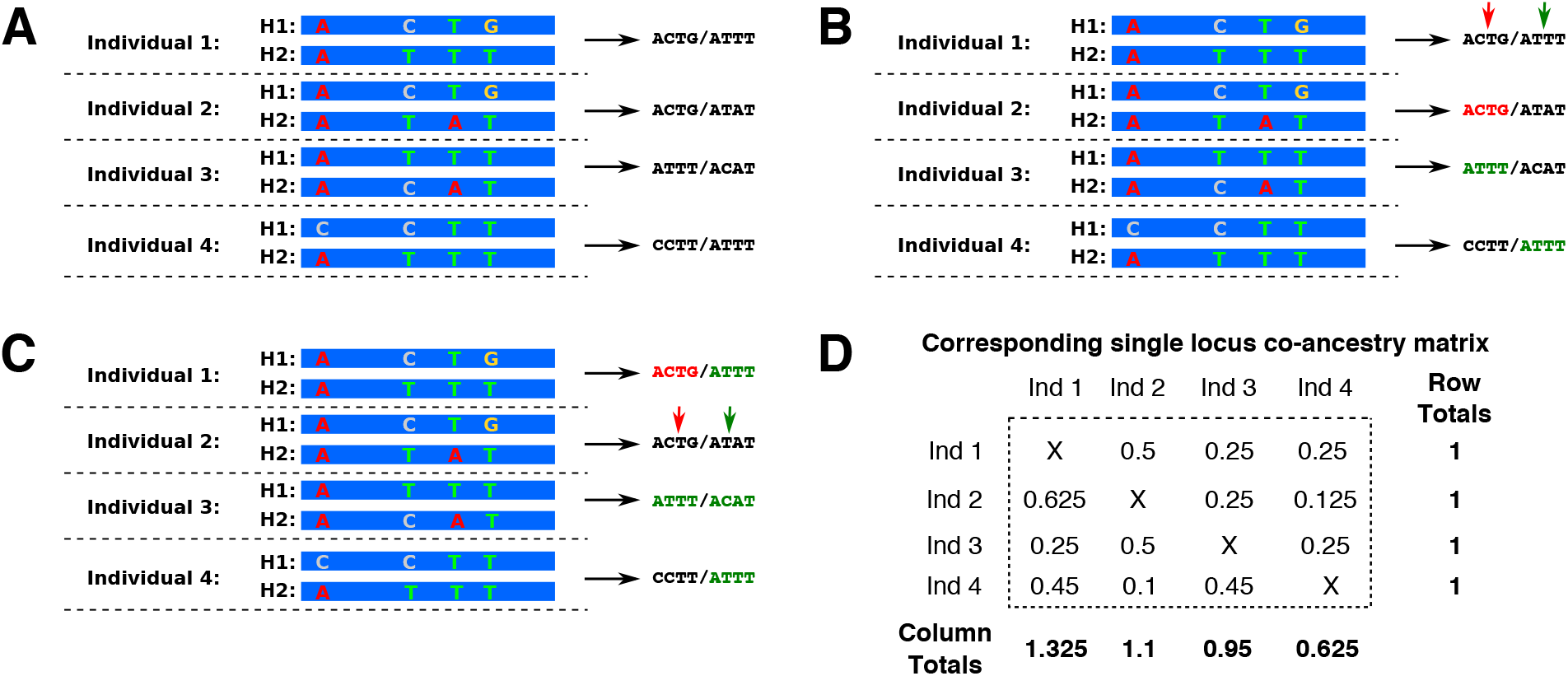
Coancestry matrix example with four diploid individuals and no missing data. **(A)** For each RAD locus (stack), we use all the variable sites in the stack to define the stack haplotypes H1 and H2. The calculation proceeds by finding the closest relatives for each haplotype. **(B)** With individual 1 as the recipient (filling the first row of the coancestry matrix), we find that the single closest relative of H1 in individual 1 is the haplotype H1 in individual 2. These haplotypes are identical (ACTG; red). Haplotype H2 in individual 1 has two closest relatives, both with an identical sequence (ATTT; green). **(C)** Focusing on individual 2 as the recipient (filling the second row of the coancestry matrix) we find that haplotype H1 in individual 2 has a single closest relative: H1 in individual 1, in a mirror image of (B). However, H2 in individual 2 does not have an identical closest relative. Its ‘donors’ are the four haplotypes that differ from it by a single SNP. **(D)** The coancestry matrix corresponding to the nearest-neighbour haplotype relationships at this locus. Note that the matrix is not symmetrical: row totals are identical for each individual and correspond to the number of RAD loci analysed. In contrast, column totals vary. Individuals with ‘outlier’ sequences are the least likely donors, CCTT in individual 4 being a clear outlier haplotype.

Concatenating all the variable sites (SNPs) at each locus to define alleles increases the resolution of `RADpainter` when compared with methods that only use a single SNP from each locus. This can be seen clearly by considering the fact that a single biallelic SNP splits the alleles at a locus into only two groups, whereas multiple linked SNPs typically enable more refined assignment of nearest neighbour relationships, as illustrated in Figure 1. Given the short length of each RAD locus (typically <250bp), we assume that all SNPs within the locus are in almost complete linkage disequilibrium (i.e. that D’ ≈ 1) and, therefore, historical recombination can be disregarded.

On the other hand, the summation across loci assumes frequent recombination between different RAD sequences - each RAD locus is counted as if providing independent evidence with regards to coancestry. This assumption is not always realistic, in particular when the dataset contains pairs of loci from the two sides of each restriction site. However, we account for any linkage between RAD loci by adjusting the normalising constant *c* which is passed on to the fineSTRUCTURE clustering algorithm together with the co-ancestry matrix and determines its sensitivity (Lawson et al. 2012).

Briefly, the `fineSTRUCTURE` clustering algorithm (Lawson et al. 2012) uses a MCMC scheme to explore the space of population configurations (sample ‘partitions’) by proposing merging or splitting populations, merging then re-splitting, or moving individuals. A proposed population configuration is accepted with a probability derived from the ratio of the likelihood with the previous configuration, a likelihood that in turn depends on the terms of the coancestry matrix scaled by the c value. Given the final `fineSTRUCTURE` output, we can infer the number of clusters, deal with a very large number of potential clusters, quantify ancestry sources in each group, and also explore relationships between groups, including with a simple tree-building algorithm (Lawson et al. 2012). Our new `fineRADstructure` package includes a set of R scripts that can be used to plot the results, including the clustered coancestry matrix, the tree with posterior population assignment probabilities, and a matrix with coancestry averages per population.

As in the original `chromoPainter` tool, the aim of estimating the c parameter is to correct for the true underlying variance of the entries of the coancestry matrix (*x_ij_*), so that the multinomial likelihood used in the clustering step matches the true statistical uncertainty in the matrix. In order to estimate an appropriate *c* value for each dataset, `RADpainter` calculates empirical variance (*V_E_*) for each entry of the coancestry matrix by jackknife, by default in blocks of 100 consecutive RAD loci. The *V_E_* entries are divided by the theoretical variance under the multinomial distribution for the coancestry matrix, i.e. assuming that an element *x_ij_* follows the binomial distribution: *x_ij_*~Binomial(*N,P_ij_*) where *N* is the total number of RAD loci in the dataset and *P_ij_* is the probability of individual *j* being the closest relative of individual *i* at any particular locus.

If linkage disequilibrium (LD) between RAD loci is weak or if the data have been mapped and sorted according to genome coordinates (so that loci in LD are grouped together within the jackknife blocks), then `RADpainter` correctly estimates the effective number of independent loci. However, when loci are not sorted and LD between them is strong then the estimation procedure may underestimate c and so be overconfident in population splits. To combat this we provide a script which reorders the RAD loci according to LD. We recommend users with unmapped data to run this before the `RADpainter` procedure in order to ensure they obtain a conservative upper bound on the number of statistically identifiable clusters. To test this approach, we constructed a RAD dataset with pairs of loci in perfect LD by duplicating all loci in a RAD dataset and randomly shuffling the positions of the duplicates. We found that *c* doubled after LD-based reordering of loci, estimating the same number of effectively independent loci correctly.

Unknown nucleotides (Ns) can be present within alleles and their positions are ignored in all pairwise sequence comparisons. Where the entire donor and/or the recipient alleles are missing, their coancestry is assumed to be proportional to the amount of coancestry observed between them in the rest of the data (i.e. ‘missing data’ coancestry is shared proportionally to ‘observed data’ coancestry). However, it is well known that one of the causes of missing alleles in RAD data is the presence of genetic polymorphisms in the restriction sites, referred to as allele dropout. Because allele dropout can lead to non-random missingness and thus influence inferences of population divergence (Arnold et al. 2013; Gautier et al. 2013), we suggest caution. The most problematic loci may be removed by filtering loci with a large excess of null alleles. In addition, we recommend that users should check for occurrence of large systematic differences in missingness between putative populations, and urge caution in interpreting the results if such differences occur. In some cases it may be appropriate to remove outlier individuals with large amount of missingness prior to a rerun of the analysis pipeline. To assist these steps, each run of `RADpainter` outputs missingness (the proportion of missing alleles) per individual.

To facilitate the use of `fineRADstructure` with existing RAD-seq processing pipelines, the input file can be generated by the widely used Stacks RAD-seq tool set (Catchen et al. 2013). This can be done directly, via output from the `export_sql.pl` Stacks program. In addition, for users who do not use the Stacks SQL database, we provide a data conversion and filtering script `Stacks2fineRAD.py` for processing the output from the core Stacks `populations` program. A third party utility script (https://github.com/edgardomortiz/fineRADstructure-tools) also enables conversion from the format of the pyRAD and ipyrad toolkits (Eaton 2014). Finally, our input format is a simple flat text file and we provide an example dataset to enable the users of other pipelines to prepare their data.

## Results

We applied `fineRADstructure` to a single-digestion RAD dataset including 120 individuals from 12 populations of the alpine plant species complex *Heliosperma pusillum*. The dataset comprises 1,097 loci which have been assembled through the *Stacks* pipeline without a reference genome (Trucchi et al. 2017). The complicated network of relationships among these twelve populations belonging to two phylogenetically intertwined species (*H. pusillum*: P, *H. veselskyi*: V), with contrasting ecology and a post-glacial history of divergence in some of the six sampled localities (A to F; Figure 2), make it an excellent case to study the performance of our approach.

The `fineRADstructure` results (a clustered coancestry matrix; Figure 2A) make the presence of twelve populations immediately clear, with substructure suggested in some of the populations. The relationships between some of the populations (A, B, C and D) are clearly not tree-like with strong evidence of heterogeneous gene flow between the species (Trucchi et al. 2017). A variable level of intra-population co-ancestry, likely related to different degree of isolation is also visible across the populations (e.g. the two populations sampled at locality F are highly isolated from each other and also the most distinct from populations sampled at all the other localities).

A number of benefits of our method can be seen in comparison against other approaches commonly used for population inference. In particular, the analysis using fastSTRUCTURE (Raj et al. 2014) at K=6 supported a clustering mainly by locality, with the exception of populations in B and C which clustered by ecology (Figure 2B). The choice of model complexity K=6 was based on the rate of decrease in the value of the Bayesian Information Criterion (BIC) (Figure 2B) as returned by the *find.clusters* function in the *R* package *adegenet* (Jombart and Ahmed 2011) and follows the methodology used in the original manuscript that presented the data (Trucchi et al. 2017). Principal Component Analysis (PCA) implemented by the glPCA function in adegenet (Jombart and Ahmed 2011) was also unable to clearly partition the genetic diversity in twelve populations (Figure 2C). The genetic structure was partly resolved by PCA by re-running the analysis including only samples from populations in localities A, B, C, D (Figure 2C; inset), but substantial overlaps remained between the population clusters, thus still providing lower resolution than `fineRADstructure`.

Overall, in terms of biological insight, a major improvement of our approach over previous results in this dataset is the clear visibility, at the same time, of both a global structure produced by a post-glacial history of recolonisation of newly ice-free mountain areas and a local structure related to parallel ecological divergence; previously these two phenomena were inferred only by applying a combination of multiple different analyses (Trucchi et al. 2017).

**Figure 2.**
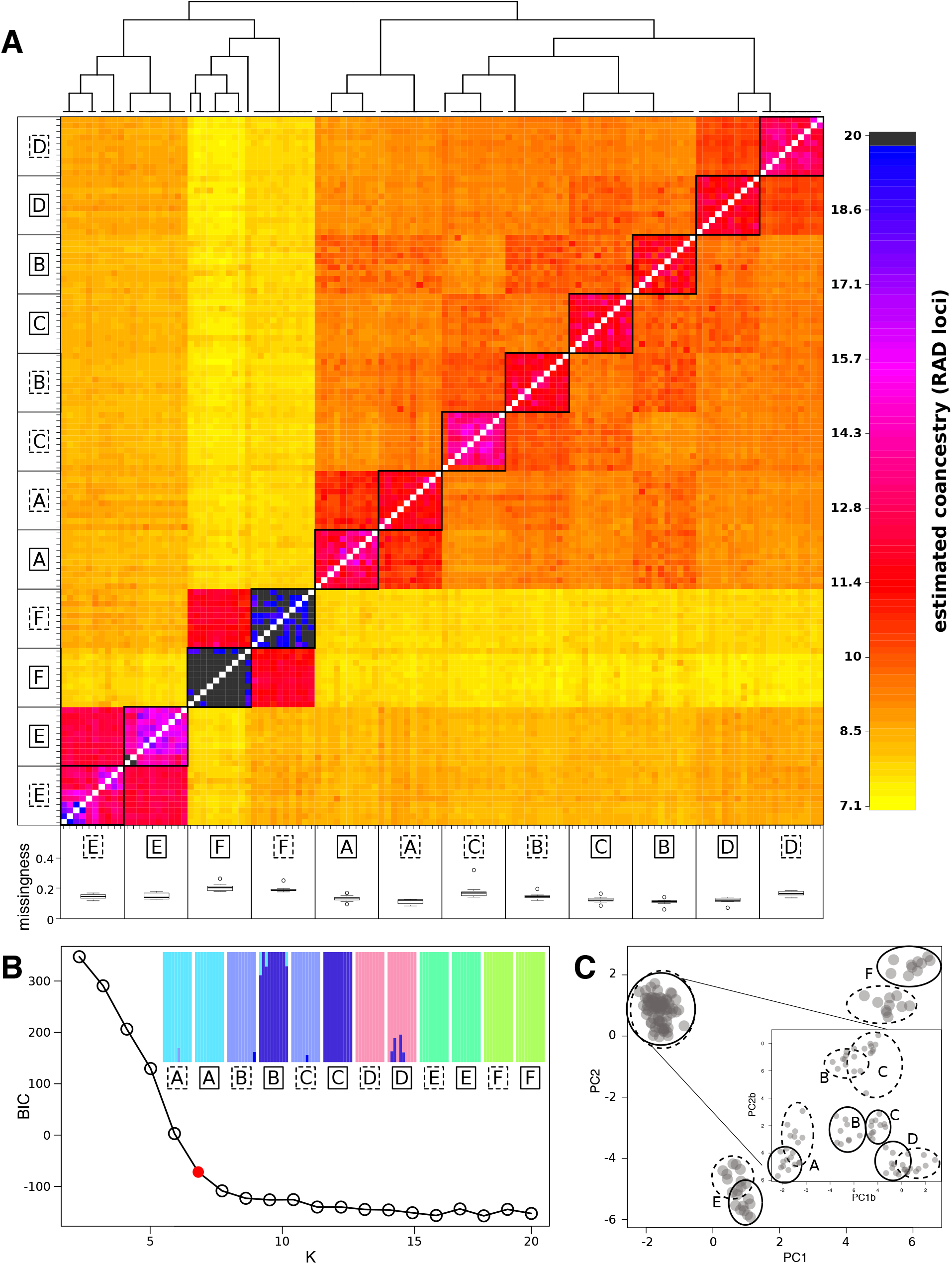
Method comparison: fineRADstructure, fastSTRUCTURE, and adegenet PCA. The analyses include individuals from six localities (A to F) and two sibling species (*H. pusillum*: solid line, *H. veselskyi*: dashed line). (A) Clustered `fineRADstructure` coancestry matrix. Individuals within the twelve populations share more coancestry with each other than between populations. Hierarchical structure among and between localities is clearly inferred-populations at localities B and C cluster by species, whereas populations at localities A, D, E, and F cluster by locality. Missingness is shown below each population label, confirming that it does not vary systematically between inferred populations. (B) fastSTRUCTURE results are shown at the suggested model complexity K=6; the inflection point where the goodness-of-fit curve - BIC estimated for increasing number of K changes its slope. (C) PCA as implemented by the adegenet package. The main plot uses all samples, while the inset shows PCA performed only on individuals from population pairs A–D.

Next we used a test dataset produced using the double digest RAD (ddRAD) approach (Peterson et al. 2012) to assess the robustness of `fineRADstructure` to non-random data missingness, specifically to the batch effect commonly present especially in ddRAD datasets, and to allele dropout. The dataset includes 76 samples from two different sub-species and three hybrid individuals. The ddRAD protocol is very sensitive to inconsistent size selection across different libraries as shifts in the size range directly influences which loci are included and sequenced in each library. Five different libraries (l1-l5), each of which underwent size selection (on a ThermoFischer E-gel) and sequencing on a PGM Ion torrent machine separately, are included in this dataset. We pre-processed the data using the common Stacks RADseq pipeline (Catchen et al. 2013). Briefly, after trimming to 200 bp, raw reads were de-multiplexed and quality filtered using the `process_radtags.pl` program, RAD loci were then assembled without a reference genome and SNPs called using the `denovo_map.pl` program (parameters `-m 5, -M 6, -n 8`) and, finally, the `populations` program was used to filter and export the loci (parameters `-r 0.8, -p 1, -min_maf 0.05 -vcf_haplotypes`).

We the used our script `StacksPopulationHaplotypes2fineRADpainter.py` (a part of our software package) to remove samples with more than 20% missing loci (Figure 3A) and loci with more than five SNPs. To assess the effect of missing data, our script transforms the resulting genotype matrix into a presence/absence matrix (replacing any genotype with 1 and missing data with 0) and produces a PCA ordination of the samples (`sklearn.decomposition` package in python).

To evaluate the effect of systematic missingness biases on `fineRADstructure` results, we compared the ‘missingness’ PCA with the co-ancestry matrix (Figure 3B,C). This comparison clearly shows that non-random missingness due to batch (i.e. library) effect is not at all mirrored in the genetic structure inferred by `fineRADstructure`; individual samples from the different sequencing libraries are intermixed within the inferred subspecies clusters (Figure 3C). Additionally, genetic hybrids between the two subspecies are clearly identified by `fineRADstructure` (as having more equal levels of coancestry with both of the subspecies; Figure 3C), despite their missing data profile being more similar to subspecies1. Thus, despite considerable non-random missingness due to genetic divergence (i.e. allele dropout), the true genetic relationship is reflected in the coancestry matrix and the missingness signal does not dominate.

**Figure 3.**
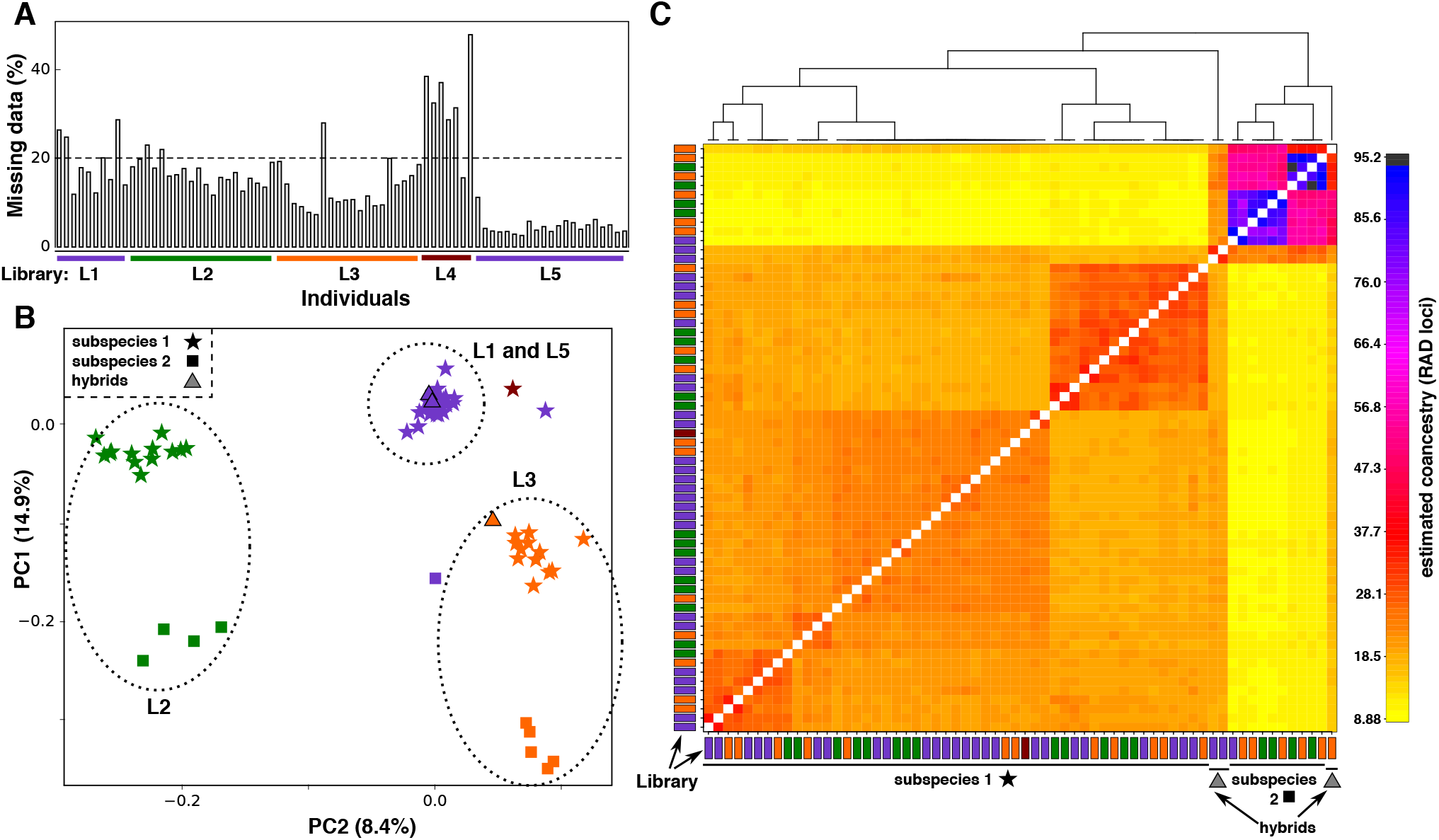
Assessing the robustness of fineRADstructure inference to systematic missingness differences. (**A**) Per sample missingness in the original dataset. We removed all individuals with missingness >20% (dashed line). **(B)** A ‘missingness PCA’, based on allele presence/absence matrix. Samples clearly cluster by library (L1 to L5; identified by colour as in (A)) due to batch effect and by sub-species due to allele dropout. **(C)** Clustered `fineRADstructure` coancestry matrix. Individuals from the two subspecies cluster together, with further sub-structure visible within the subspecies. Importantly, the structure within the subspecies does not relate to the missingness PCA clusters (i.e. it does not relate to the library batch). Hybrid individuals are clearly identifiable, sharing relatively more equal coancestry levels with both subspecies.

## Discussion

In this manuscript, we have described software that enables fine population structure inference based on nearest neighbour (first coalescence) relationship between haplotypes inferred from RAD-seq data. Thus, our software brings the benefits of this approach especially to genomics of non-model organisms, which currently relies heavily on RAD-seq. In this context, it is useful to emphasise two considerations related to the use of `fineRADstructure` across a range of species.

First, because we assume independence between RAD loci, the coancestry matrix entries do not depend on RAD loci being ordered along the genome. Therefore, analyses with and without a reference genome will produce identical coancestry matrices. Second, our method benefits from linkage (LD) between multiple polymorphisms within the same RAD locus, enabling a more specific assignment of nearest neighbour relationships between individuals, as shown in Figure 1. Therefore, unlike many methods which assume markers are unlinked and require removal of polymorphisms that are in high LD with each other (e.g. PCA based methods, STRUCTURE), `fineRADstructure` gains additional power from their presence within a RAD locus, as will be the case especially in species with high nucleotide polymorphism levels.

`fineRADstructure` aims to detect and report all excess sample relatedness that rises above statistical noise in the coancestry matrix, i.e. a population is defined as a group of individuals with indistinguishable genetic ancestry in the dataset. This approach resulted in biologically meaningful clustering for all the datasets we tested. However, we would caution against interpreting the number of clusters as a definite estimate of the ‘correct’ number of populations. As seen for example in Figure 2A, species often contain different levels of population sub-structure. Therefore, we suggests that especially in complex datasets users may sometimes obtain a better intuition about the relationships between their samples by repeating the clustering step several times with varying sensitivity (*c* parameter values) and interpreting the results jointly.

Finally, we would like to point out that although the *ad hoc* tree building approach generally performs well, it is intended to be illustrative of the relationships between the populations, rather than to represent the true population history. Therefore, we suggests that where the true historical relationships are known (e.g. as a result of independent phylogenetic analysis), the clustering step can be skipped and the coancestry matrix ordered according to the pre-existing information. Under those circumstances, the entries of the coancestry matrix provide an independent source of information on recent sample relatedness, complementing the information about older historical relationships; this may be especially informative in cases of recent gene-flow between populations.

## Acknowledgements

This work was supported by the Medical Research Council (MR/M501608/1 to D.F), ), the Austrian Climate Research Programme (ACRP5-EpiChange-KR12AC5K01286), the Austrian Science Fund (FWF, Y661-B16) for E.T., and the Wellcome Trust (097677/Z/11/Z to M.M.). The idea for `RADpainter` and `fineRADstructure` arose in discussions at the 2016 Workshop on Population and Speciation Genomics in Cesky Krumlov (http://evomics.org); we would like to thank all the faculty and participants in the workshop.

